# Finite Element and deformation analyses predict pattern of bone failure in loaded zebrafish spines

**DOI:** 10.1101/703629

**Authors:** Elis Newman, Erika Kague, Jessye A. Aggleton, Christianne Fernee, Kate Robson Brown, Chrissy L Hammond

## Abstract

The spine is the central skeletal support structure in vertebrates consisting of repeated units of bone, the vertebrae, separated by intervertebral discs that enable the movement of the spine. Spinal pathologies such as idiopathic back pain, vertebral compression fractures and intervertebral disc failure affect millions of people world-wide. Animal models can help us to understand the disease process, and zebrafish are increasingly used as they are highly genetically tractable, their spines are axially loaded like humans, and they show similar pathologies to humans during ageing. However biomechanical models for the zebrafish are largely lacking. Here we describe the results of loading intact zebrafish spinal motion segments on a material testing stage within a micro Computed Tomography machine. We show that vertebrae and their arches show predictable patterns of deformation prior to their ultimate failure, in a pattern dependent on their position within the segment. We further show using geometric morphometrics which regions of the vertebra deform the most during loading, and that Finite Element models of the trunk subjected reflect the real patterns of deformation and strain seen during loading and can therefore be used as a predictive model for biomechanical performance.

## Introduction

The spine consists of a repeated pattern of motion segments (MSs) of bony vertebrae separated by intervertebral disks (IVDs) that enable spinal movement. Back pain and intervertebral disc degeneration affect hundreds of millions of people worldwide (1,2), and vertebral compression fractures are a frequent feature of osteoporosis (3). Biomechanical pathologies of the spine are underpinned by genetic, physiological and environmental pathways that together damage the soft tissues (IVD and muscle) and/or the bone changing the mechanics of the system.

Animal models, typically rodents, are frequently used to study mechanisms of spinal pathology (4). However, quadrupeds are disadvantageous for studying the human spine as gravitational load acts perpendicular to their axial skeleton. Zebrafish are increasingly used as a model for human disease, due to their genetic tractability. Unlike quadrupeds, their spine is axially loaded as a result of swimming through viscose water (5). Zebrafish are well established as models for skeletogenesis, pathology, and ageing (6), and develop spinal pathologies in response to altered genetics (7) and ageing (8). However, the biomechanics of the zebrafish spine are comparatively poorly characterised.

Finite element analysis (FEA) has proven a pivotal tool in the study of biomechanical subjects (9), and offers a method for biomechanically characterising the zebrafish spine, including intact motion segments. This technique digitally models an object of known material properties using a series of linked nodes of known number and geometry, that can be subjected to a wide variety of forces outputting the predicted geometry, strain and deformation. Results can be validated by comparison with the results of loading experiments in which a sample is loaded *ex vivo* (10,11). FEA has been used in zebrafish to identify regions of high strain and test contributions of shape and material properties in joint morphogenesis (12,13) and to study strain patterns in a single vertebra (14).

Here, we describe a novel integrated experimental platform that brings together imaging, modelling and real-world loading validation to explore the biomechanics of intact zebrafish spinal motion segments. We generated an FEA model of the spine, which we validated with a loading experiment using a high-precision material testing stage (MTS) under set loading regimes using micro Computed Tomography (μ-CT). Three-dimensional geometric morphometrics (3D GM) was used to explore the patterns of deformation seen in each vertebra during loading. Comparison of results demonstrated that our FEA model accurately predicted the relative patterns of deformation and strain experienced by real samples loaded *ex vivo.*

## Methods

### Zebrafish samples

1-year old, wild-type (WT) zebrafish were fixed in 4% paraformaldehyde and dehydrated to 70% EtOH. Whole-mount MSs were acquired by making two cuts in the trunk, between vertebra 18 and 24 (Fig. 1a-c).

**Figure 1.**
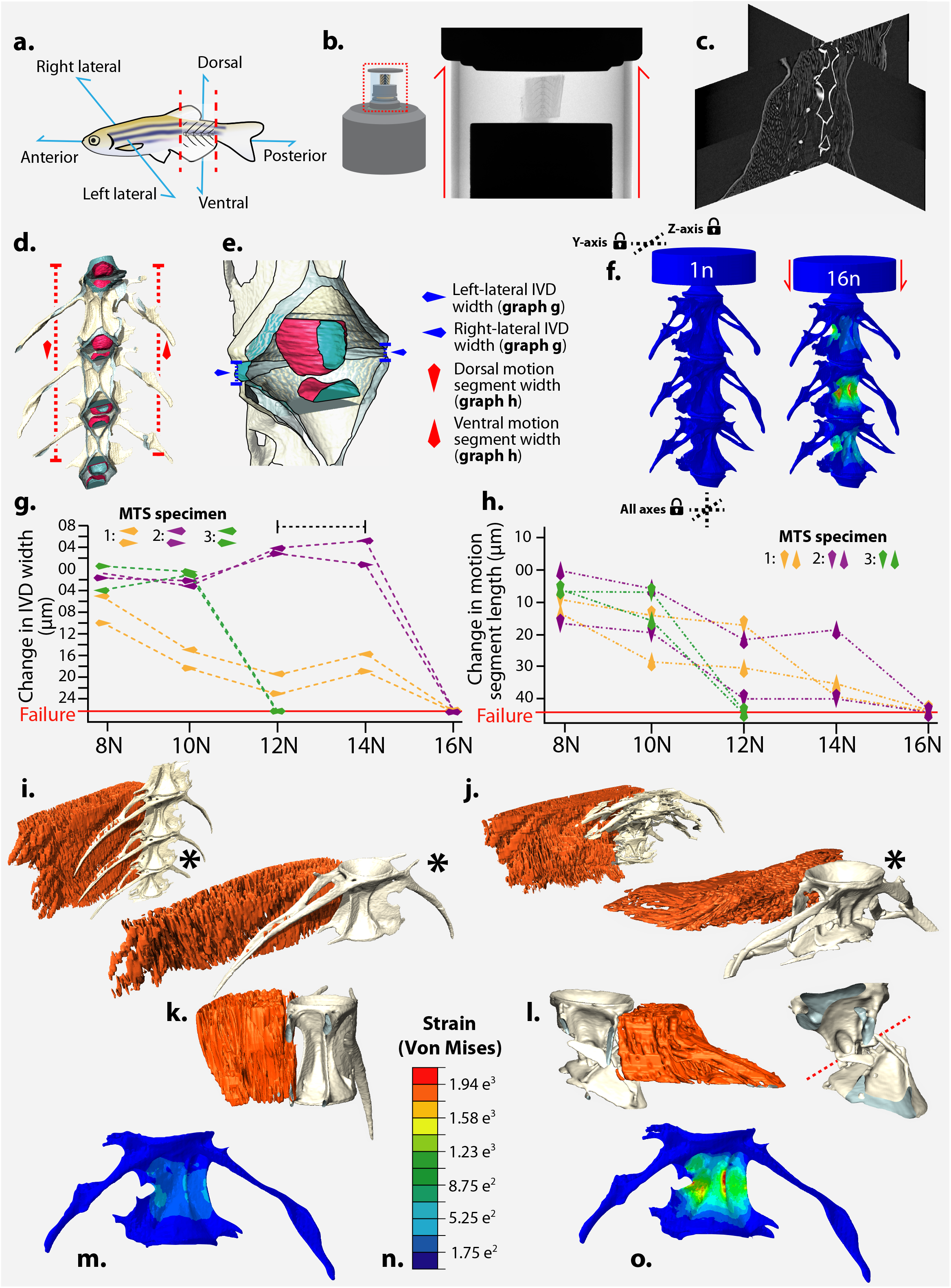
Ex vivo spine loading leads to motion segment failure in a region of high strain predicted by Finite Element Analysis. **a**, Schematic of zebrafish motion segment (MS) dissection. **b**, Material testing stage (MTS) schematic and X-radiograph. **c**, Orthogonal reconstruction slices showing vertebrae and associated soft tissue. **d**, Three-dimensional reconstruction of the finite element analysis (FEA) model with colours reflecting different materials. **E**, Details of the nucleus-pulposus (pink) and annulus fibrosis (blue) from **d** showing linear measurements of inter-vertebral disk (IVD) thickness. **f**, Predicted compressive deformation and strain map from FEA; dashed lines indicate axes in which boundary conditions were established. **g**, Changes to IVD width measurements (bracketed dashed line highlights IVD elastic rebound) and **h**, changes in MS length with increasing load for the three MTS specimens; direction of arrowhead denotes measurement type. **i, j** Reconstructions of MTS Specimen 1 compressed to 10N (**i**), and 16N (**j**) with central vertebra indicated by * in each. **K & l** Antero-posterior cross-sections of the central vertebra at 10N (**k**) and 16N (**l**). **m**, **o** FEA strain maps at 10n (**m**) and 16n (**o**). Scale shown in **n**.

### *In vitro* vertebral loading experiment

Loading experiments were conducted using a custom-built Material Testing Stage (MTS2) in the Bruker SKYSCAN 1272 μCT system. Radiographic visualisation of each MS (n=3) was carried out and if required, vertebrae were trimmed to retain three complete vertebrae (between V18 and V24) and associated IVDs (Fig. 1b-d). Samples were stabilized (anterior-up) in the MTS2 using cyanoacrylate glue. The MTS2 was programmed to perform a sequential series of seven μCT scans at a series of increasing loads (Table 1), using X-ray energy of 60 KeV, current of 50 W, isotropic voxel size of 5 μm and a 0.25 mm Aluminium filter. 1501 projections were collected during a 180° rotation, with 400 ms exposure time. Reconstructions were performed using NRecon (Version 1.7.1.0). Surfaces of vertebrae, muscle and IVDs in each dataset were generated using Avizo (Avizo version 8; Vizualisation Sciences Group)(Fig. 1c-e, Table 1) and linear measurements of IVDs and MS lengths made using the “3D Measurement” tool. Vertebrae surfaces were further processed in Meshlab (Table 2).

**Table 1.** Scanning schedule for specimens analysed using the material testing stage (MTS), and their linear morphometric measurements and Principal components Analysis (PCA) scores of landmark deformation data measured using three-dimensional geometric morphometrics (3D-GM).

| Specimen | Compressive load (N) | Left lateral motion segment length (µm) | Right lateral motion segment length (µm) | Linear inter-vertebral disk (IVD) proxy measurements (µm) |  |  |  |  |  |  |  | 3D-GM PCA scores |  |  |  |  |  |
| --- | --- | --- | --- | --- | --- | --- | --- | --- | --- | --- | --- | --- | --- | --- | --- | --- | --- |
|  |  |  |  | IVD between top and middle vertebra |  |  |  | IVD between middle and bottom vertebra |  |  |  | Anterior vertebra |  | Central vertebra |  | Posterior vertebra |  |
|  |  |  |  | Left-lateral central distance | Right-lateral central distance | Ventral central distance | Dorsal central distance | Left-lateral central distance | Right-lateral central distance | Ventral central distance | Dorsal central distance | PC1 (antero-posterior deformation) | PC2 (lateral deformation) | PC1 (antero-posterior deformation) | PC2 (lateral deformation) | PC1 (antero-posterior deformation) | PC2 (lateral deformation) |
| MTS 1 | 1 | 2116 | 2128 | 37.8 | 39.2 | 12.15 | 20.72 | 74.42 | 40.8 | 15.43 | 8.94 | -0.019 | 0.003 | -0.011 | 0.001 | -0.021 | 0.008 |
|  | 5 | 2113 | 2122 | 36.97 | 37.15 | 10.39 | 19.17 | 65.38 | 36.72 | 12.37 | 6.43 | -0.015 | 0.008 | -0.006 | -0.009 | -0.017 | 0.004 |
|  | 8 | 2104 | 2109 | 34.17 | 31.75 | 7.51 | 19.89 | 55.51 | 31.67 | 0 | 4.26 | -0.002 | -0.003 | 0.000 | 0.000 | -0.006 | -0.002 |
|  | 10 | 2099 | 2094 | 31.13 | 31.99 | 2.95 | 14.05 | 46.95 | 21.43 | 0 | 4.43 | -0.001 | -0.008 | 0.002 | 0.012 | -0.003 | -0.003 |
|  | 12 | 2096 | 2092 | 29.42 | 22.42 | 0 | 12.43 | 42.19 | 17.09 | 0 | 0 | 0.002 | -0.013 | 0.005 | 0.008 | 0.007 | -0.010 |
|  | 14 | 2074 | 2085 | 31.47 | 25.82 | 0 | 18.54 | 46.37 | 21.05 | 0 | 0 | 0.034 | 0.005 | 0.014 | -0.004 | 0.013 | 0.009 |
|  | 16 | FAILURE | FAILURE | FAILURE | FAILURE | FAILURE | FAILURE | FAILURE | FAILURE | FAILURE | FAILURE | FAILURE | FAILURE | FAILURE | FAILURE | FAILURE | FAILURE |
| MTS 2 | 1 | 2364 | 2346 | 24.65 | 13.86 | 6.8 | 10.4 | 17.25 | 15.38 | 3.2 | 6.85 | -0.026 | 0.002 | -0.014 | 0.002 | -0.015 | 0.001 |
|  | 5 | 2356 | 2344 | 21.04 | 12.56 | 5.72 | 9.98 | 16.73 | 14.12 | 0 | 0 | -0.002 | -0.006 | -0.005 | -0.002 | -0.013 | 0.001 |
|  | 8 | 2339 | 2344 | 18.21 | 12.21 | 0 | 0 | 15.04 | 13.53 | 0 | 0 | 0.002 | -0.008 | -0.001 | 0.011 | -0.010 | 0.002 |
|  | 10 | 2336 | 2338 | 16.37 | 9.56 | 0 | 0 | 14.54 | 10.93 | 0 | 0 | 0.006 | -0.008 | 0.001 | 0.005 | 0.005 | -0.015 |
|  | 12 | 2316 | 2323 | 17.2 | 12.51 | 9.76 | 7.82 | 19.55 | 17.99 | 4.5 | 7.47 | 0.006 | 0.008 | 0.002 | 0.000 | 0.012 | -0.006 |
|  | 14 | 2316 | 2326 | 18.94 | 17.53 | 11.92 | 9.4 | 17.52 | 19.32 | 9.5 | 12.34 | 0.011 | 0.004 | 0.004 | -0.004 | 0.015 | 0.008 |
|  | 16 | FAILURE | FAILURE | FAILURE | FAILURE | FAILURE | FAILURE | FAILURE | FAILURE | FAILURE | FAILURE | FAILURE | FAILURE | FAILURE | FAILURE | FAILURE | FAILURE |
| MTS 3 | 1 | 1832 | 1829 | 15.54 | 11.67 | 0 | 0 |  | 15.15 | 0 | 0 | -0.013 | 0.002 | -0.012 | 0.007 | -0.005 | -0.010 |
|  | 5 | 1827 | 1822 | 14.23 | 10.28 | 0 | 0 | 15.39 | 14.2 | 0 | 0 | 0.001 | -0.010 | -0.001 | 0.000 | 0.007 | -0.006 |
|  | 8 | 1820 | 1816 | 13.97 | 7.75 | 0 | 0 | 15.78 | 10.26 | 0 | 0 | 0.005 | 0.010 | 0.001 | -0.012 | 0.013 | 0.005 |
|  | 10 | 1821 | 1807 | 13.31 | 7.82 | 0 | 0 | 14.97 | 13.15 | 0 | 0 | 0.007 | -0.002 | 0.013 | 0.005 | 0.019 | 0.008 |
|  | 12 | FAILURE | FAILURE | FAILURE | FAILURE | FAILURE | FAILURE | FAILURE | FAILURE | FAILURE | FAILURE | FAILURE | FAILURE | FAILURE | FAILURE | FAILURE | FAILURE |
|  | 14 | FAILURE | FAILURE | FAILURE | FAILURE | FAILURE | FAILURE | FAILURE | FAILURE | FAILURE | FAILURE | FAILURE | FAILURE | FAILURE | FAILURE | FAILURE | FAILURE |
|  | 16 | FAILURE | FAILURE | FAILURE | FAILURE | FAILURE | FAILURE | FAILURE | FAILURE | FAILURE | FAILURE | FAILURE | FAILURE | FAILURE | FAILURE | FAILURE | FAILURE |

**Table 2.**
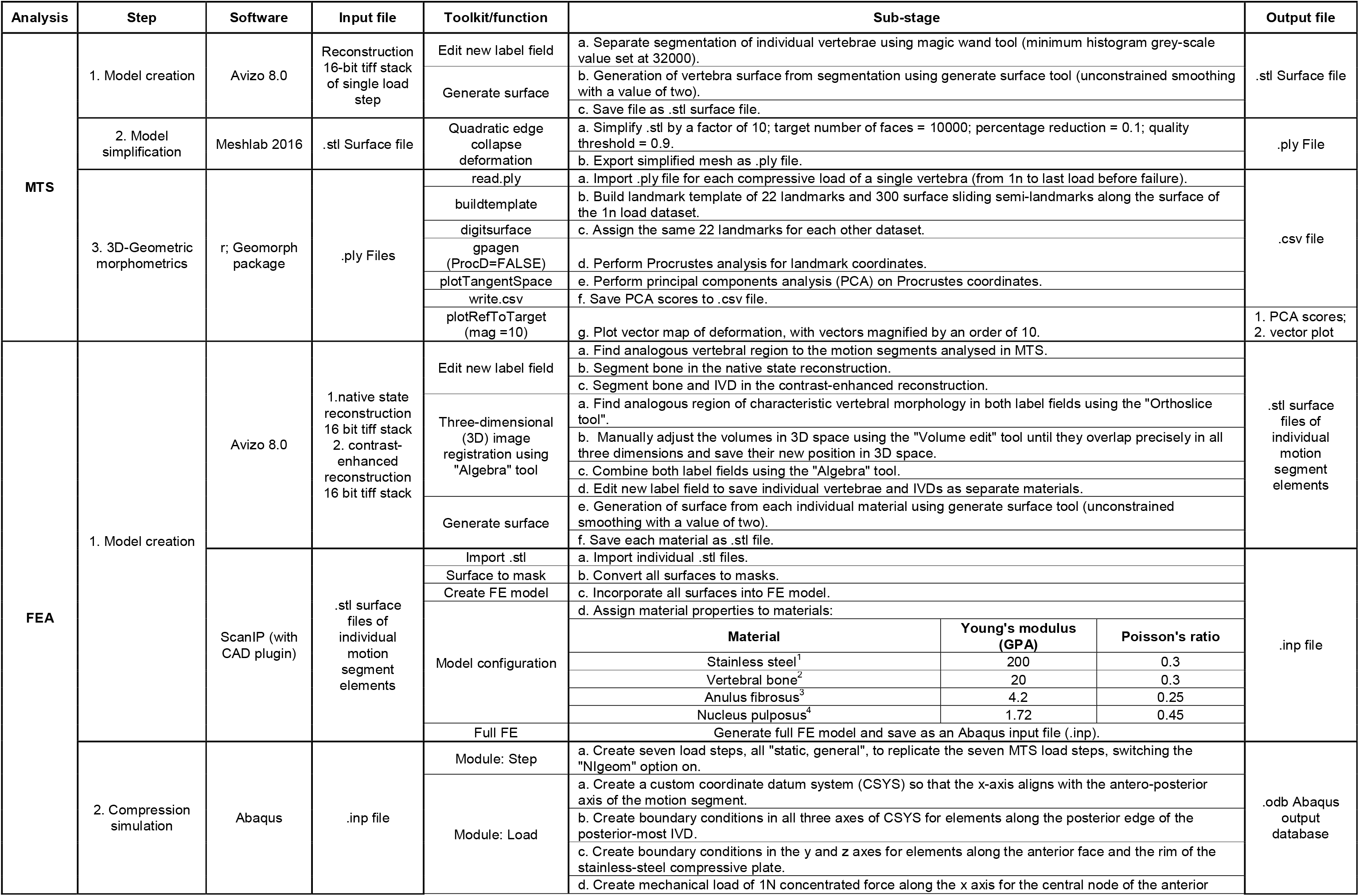
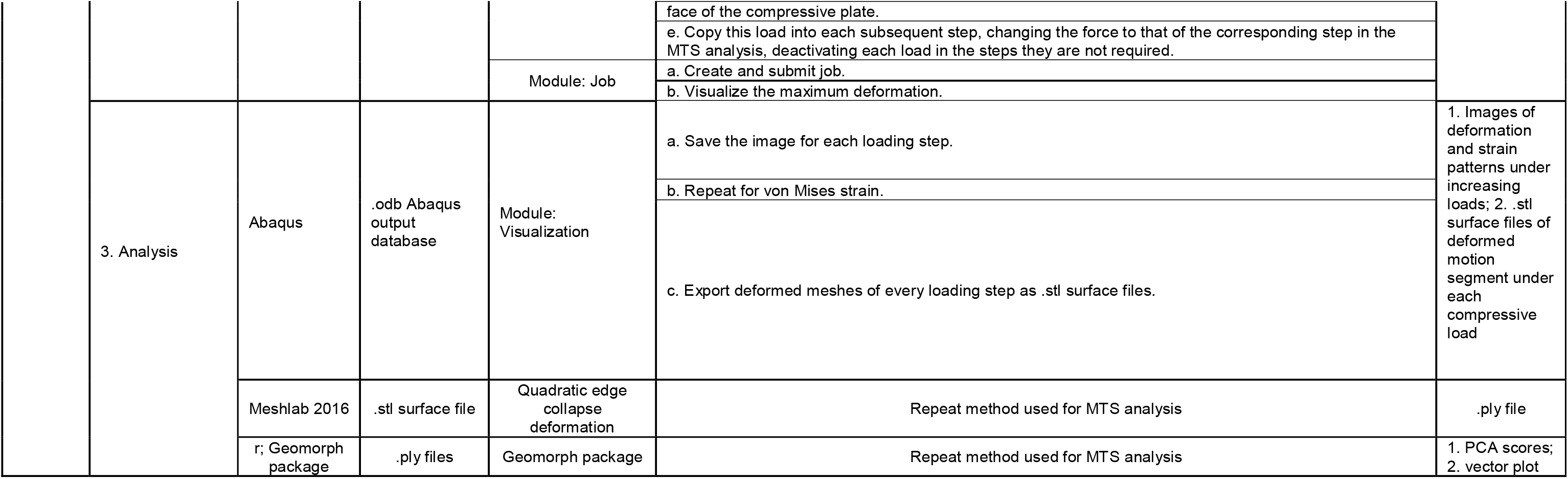
Methodology tree for image processing and analysis. 1. Material properties from the Engineering Toolbox (www.engineeringtoolbox.com; accessed 18.06.19); 2. Material properties from Ofer et al., (2019); 3. Material properties assessed using AFM of joint cartilage (Harniman *Pers. Comm.s*); 4. Material properties from Panzer and Cronin (2009)

### Finite Element Analysis (FEA)

A vertebral surface mesh was created based on a 1-year-old WT specimen μCT scanned using a Nikon XTH 225ST μCT system as described under two conditions; (a) native state and (b) contrast-enhanced following incubation for 14 days in 2.5% phosphomolybdemic acid (15). Scan (a) was used to segment hard tissues (V18-V24), and scan (b) to segment IVDs. The resulting binary label fields from scans (a) and (b) were saved as 8-bit tiff stacks, manually registered in 3D space in Avizo (‘Trackball’ tool) and algorithmically combined (‘Algebra’ tool), creating a single volume of separate materials representing three vertebrae and four IVDs (Fig. 1d-e, Table 2). A 500 μm thick cylinder was created contacting the anterior-most IVD perpendicular to the model axis, to mimic the stainless-steel compressive plate and distribution of forces applied during loading (Fig. 1f).

The complete vertebral surface mesh was imported into Simpleware ScanIP (version 2018.12, Synopsys Inc.) to create an FE model. The model consisted of 1,054,187 linear tetrahedral elements joined at 257,392 nodes comprising four material types: vertebral bone, annulus-fibrosus, nucleus-pulposus and stainless-steel (Fig. 1d-f, Table 2). The model was analysed in Abaqus (2018 version). A custom datum coordinate system was created centred on the antero-posterior axis of the model, and a concentrated force applied to the central node of the anterior face of the compressive plate. This loading case was repeated in each of 7 steps of a multi-step analysis, with load values matching the increments applied in the MTS (Table 1). The model was constrained in two locations using boundary conditions, at the base of the posterior-most IVD (constrained in 3 axes), and at the top of the compressive plate (constrained in 2 axes, allowing movement along the model’s antero-posterior axis (Fig. 1f). Deformed meshes from each step were exported as surface files and analysed using 3D-GM for direct quantitative comparison between relative and absolute patterns of deformation predicted by FEA and observed in MTS data.

### Three-dimensional geometric morphometrics (3D-GM)

Geometric morphometric analysis of vertebral shape deformation was performed using the “Geomorph” package for the “r” statistics software (16). For each loading experiment, we used the first scan (1N load) to create a template of 3D coordinates for 22 fixed three-dimensional landmarks (Fig. 2a-c) linked by 300 surface sliding semi-landmarks (using the “buildtemplate” function). By assigning the same landmarks in each scan (using the “digitsurface” function), we compared the first scan with subsequent scans of the same vertebra using generalised Procrustes analysis (allowing semi-landmarks to ‘slide’ in order to remove arbitrary spacing). Resulting shape variables were subjected to principal component analysis (PCA) to identify the principal patterns of variation between scans of the same vertebra, and isolate trends in deformation with increasing compressive load.

**Figure 2.**
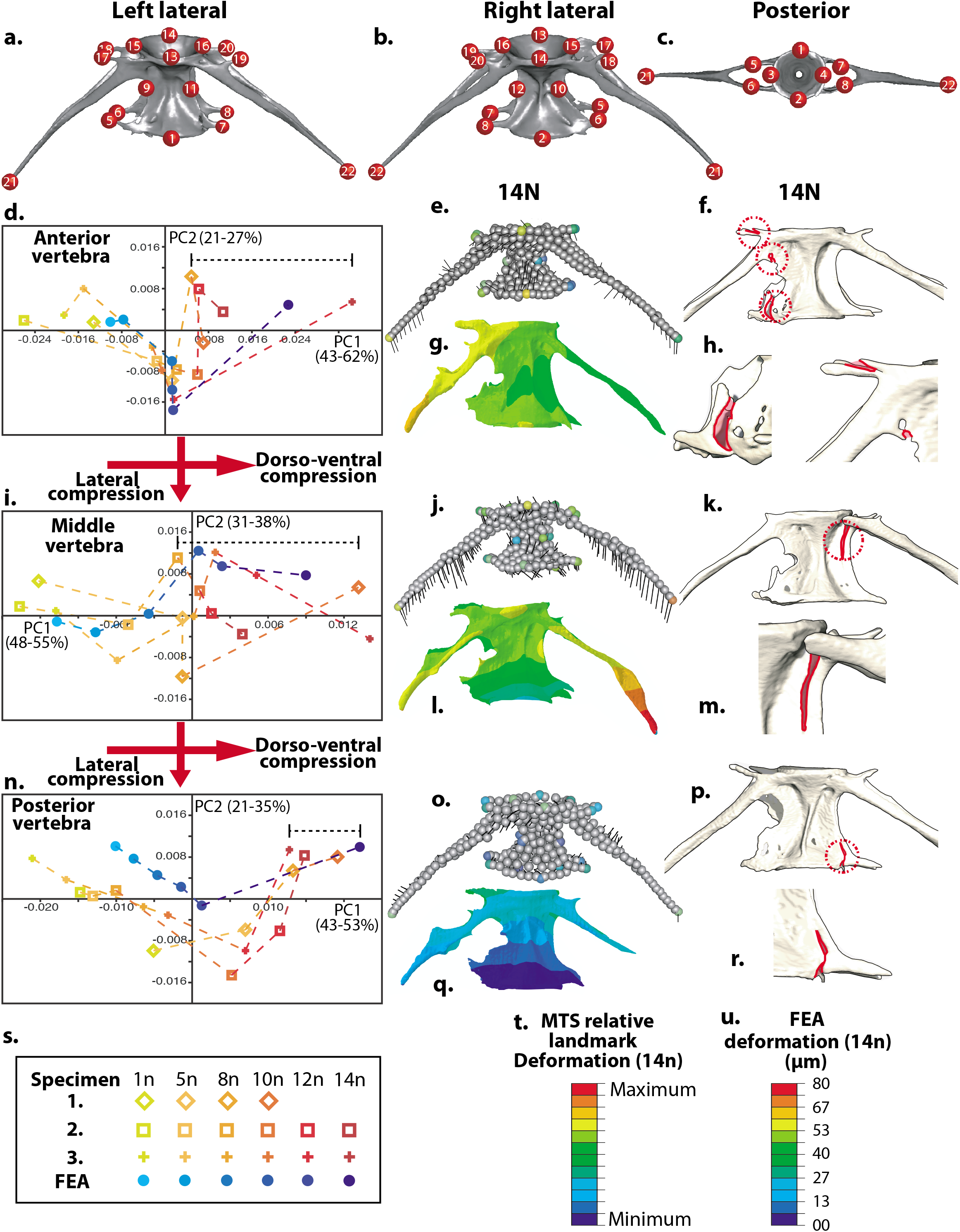
Finite Element and geometric morphometric analyses model deformation patterns prior to failure. **a-c** Landmarks assigned for Three-dimensional geometric morphometric (3D-GM) analysis. **d, i & n** Results of principal components analyses (PCA) of landmark deformation under increasing compressive loads for each specimen, and deformation predicted by FEA (key in **s**). Black bracketed lines indicate reduced lateral compression. **e, j & o** 3D vector plots with vectors and colours highlighting the direction and extent of landmark deformation for each vertebra in Specimen 1 (extents magnified by 10; colour scale in **t**). **g, i & q** Deformation maps predicted by FEA (scales presented in **u**). **f, h, k, m, p & r** Examples of fractures (outlined in red for clarity) occurring at compressive loads before failure; corresponding with deformation patterns predicted in FEA and seen *Ex vivo.*

## 3. Results and discussion

### Vertebral motion segments fail under loading of 12-16N at positions of maximum Von Mises strain

To test the range of compressive loads that the motion segment (MS) could resist until failure, we subjected an MS to exponentially increasing compressive forces from 1-100N. This specimen failed at 16N whereupon the central vertebra fractured mid-centrum. A primary loading regime between 1-16N was thus established (Table 1) for the three primary specimens; occupying the elastic, plastic and failure regions of the compressive loading profile of a typical MS. Failure was considered when at least one vertebral centrum fractured across the axis (e.g. Fig. 1j,l). All samples failed between 12-16N upon shallow angle fracture in the central vertebra. Prior to failure, linear measurements show an increase in IVD antero-posterior thickness (Table 1, Fig. 1g) suggesting elastic rebound of the IVD that may further contribute to the ultimate strain and failure of the segment (Fig. 1h). The surrounding epaxial musculature showed no obvious deformation or damage until the entire system failed upon vertebral fracture, at which point muscle fibre organisation was lost (Fig 1i-l). Comparison between MTS μCT data and the FEA results demonstrated strong spatial correlation between maximum predicted strain and ultimate point of sample failure in the central vertebra (Fig 1.m-o).

### Morphometric characterization of vertebral compression is predicted by FEA

We found characteristic patterns of deformation and strain in response to compressive loading of zebrafish vertebrae. 3D GM results from MTS data follow distinct trends for each vertebrae between the three specimens (Fig.2d,i,n), showing consistent dorso-ventral compression and lateral compression that is reversed at higher loads, potentially due to elastic rebound of the IVD and fracturing along the zygopophyses that occurs at these loads (Fig. 2). Fractures are observed where the arches and zygopophyses contact the centrum, at loads that precede the failure of the segment (Fig 2f, h, k, m, p & r). Comparison with FEA data (blue points in graphs d,i,n) suggest that the FE model accurately predicts these patterns (Fig. 2d,i & n), and that patterns of deformation could explain the first signs of damage prior to failure. In both datasets the anterior vertebra undergoes most deformation, particularly of the arches which are deformed posteriorly (Fig. 2 e-h). The central vertebrae and arches show strong torsion (Fig. j-m), increasing through the loading regime leading to the failure of the segment (Fig. 1l,o). The posterior vertebra shows the least deformation and is most isotropic in pattern (Fig. 2o-r), potentially due to protection offered by the anterior IVDs.

Comparison with *ex-vivo* loading of vertebral MSs validates the accuracy of our FEA model for predicting patterns of deformation and strain across these structures. This offers a step towards a digital ‘sandbox’ approach to modelling the effects of genetic, physiological and morphological properties on the reaction and resistance of vertebral MSs to loading. Inputting specific properties of vertebral samples into a validated FE model will allow their effects on the biomechanics of the spine to be quantitatively tested *in silico,* allowing the relative contributions of shape and material properties to be explored and empirically tested. This will aid comparison of mechanical performance between different model systems. As an advantage of the zebrafish system is the wealth of mutants modelling human disease genetics (17), comparisons of mechanical performance between genotype and phenotype will be possible.In the longer term this approach may give insight into biomechanical aspects of spinal pathology; allowing identification of ‘at risk sites’ in the spine. This could provide a basis for more specific or earlier interventions than those commonly employed.

## Acknowledgements and Funding

We would like to thank Rob Harniman for the AFM values for cartilage (acquired for another project). EN EK CLH and KRB were funded by STFC grant ST/T000678/1 and CLH and EK by Versus Arthritis Fellowship 21937 and project grant 21211

## Conflict of interest

The authors report no conflicts of interest

## Author contributions

EN, EK and JA performed experiments, EN, EK, JA, CF and CLH analysed data.

The project was designed by CH and KRB. All authors contributed to drafting the manuscript.

## Data availability

Models are available at data.bris.ac.uk

